# Genomic prediction of autotetraploids; influence of relationship matrices, allele dosage, and continuous genotyping calls in the phenotype prediction

**DOI:** 10.1101/432179

**Authors:** Ivone de Bem Oliveira, Marcio F. R. Resende, Luis Felipe V. Ferrão, Rodrigo R. Amadeu, Jeffrey B. Endelman, Matias Kirst, Alexandre S. G. Coelho, Patricio R. Munoz

**Author notes:** **Corresponding author:** Patricio Ricardo Muñoz, Blueberry Breeding & Genomics Lab, University of Florida, Address: 2550 Hull Road, P.O. Box 110690, Gainesville, FL, 32611.

## Abstract

Estimation of allele dosage in autopolyploids is challenging and current methods often result in the misclassification of genotypes. Here we propose and compare the use of next generation sequencing read depth as continuous parameterization for autotetraploid genomic prediction of breeding values, using blueberry (*Vaccinium corybosum* spp.) as a model. Additionally, we investigated the influence of different sources of information to build relationship matrices in phenotype prediction; no relationship, pedigree, and genomic information, considering either diploid or tetraploid parameterizations. A real breeding population composed of 1,847 individuals was phenotyped for eight yield and fruit quality traits over two years. Analyses were based on extensive pedigree (since 1908) and high-density marker data (86K markers). Our results show that marker-based matrices can yield significantly better prediction than pedigree for most of the traits, based on model fitting and expected genetic gain. Continuous genotypic based models performed as well as the current best models and presented a significantly better goodness-of-fit for all traits analyzed. This approach also reduces the computational time required for marker calling and avoids problems associated with misclassification of genotypic classes when assigning dosage in polyploid species. Accuracies are encouraging for application of genomic selection (GS) for blueberry breeding. Conservatively, GS could reduce the time for cultivar release by three years. GS could increase the genetic gain per cycle by 86% on average when compared to phenotypic selection, and 32% when compared with pedigree-based selection.

## INTRODUCTION

Polyploidy events are not an exception in plants, as about 70% of Angiosperms and 95% of Pteridophytes underwent at least one polyploidization event (Soltis and Soltis 1999). Polyploids are normally grouped into two categories, autopolyploids and allopolyploids, but intermediate forms are also possible, such as segmental allopolyploids (Spoelhof *et al.* 2017). Thresholds for polyploid classification have been controversial, but following the general taxonomic definition, autopolyploids arise from within-species whole genome duplication, and allopolyploids arise from whole genome duplication prior to or after an inter-specific hybridization event (Soltis *et al.* 2007).

Because speciation via ploidy increase can generate new phenotypic variability, this phenomenon is considered a powerful evolutionary source (Hieter and Griffiths 1999; Soltis *et al.* 2016). Despite the important role of polyploidization in plant evolution, its effects on inheritance of many agronomic traits and population genetics are still poorly understood when compared with diploid species (Dufresne *et al.* 2014). This especially holds true for autopolyploids. The complex nature of autopolyploid genetics is due to the presence of genotypes with higher allele dosage than diploids, larger number of genotypic classes, possibility of multivalent pairing, and poor knowledge of chromosome behavior during meiosis (Slater *et al.* 2013; Dufresne *et al.* 2014; Mollinari *et al.* 2015).

The advent of high-throughput genotyping methods, associated with the development of genetic and statistical analysis tools, has generated significant genetic gains for diploid species (Desta and Ortiz 2014). However, the application of genomic information to polyploid crops remains a challenge (Comai *et al.* 2005; Grandke *et al.* 2016). Although methods for the analysis and interpretation of genetic data in polyploids have recently been described (see review in Bourke *et al.* 2018), much development is needed, especially for new breeding approaches, such as genomic selection.

Genomic selection (GS) is a method to increase the efficiency and accelerate the selection process in breeding programs. GS is used to capture the simultaneous effects of molecular markers distributed across the genome, based in the premise that linkage disequilibrium between causal polymorphisms and markers allow the prediction of phenotypes based on the genotypic values (Meuwissen *et al.* 2001; Zhang *et al.* 2011; de los Campos *et al.* 2013). The first GS studies addressing polyploids considered diploid genetic models to circumvent the complexity involved in accurately defining allelic dosage (i.e., the number of copies of each allele at a given polymorphic locus). Promising results have been reported for polyploids (e.g. Gouy *et al.* 2013; Annicchiarico *et al.* 2015; Ashraf *et al.* 2016), however simplified assumptions were mostly used for genetic and statistical inferences (Garcia *et al.* 2013). Only a few studies have added different factors accounting for polyploid effects (*e.g.*, Slater *et al.* 2016; Sverrisdottir el al. 2017). Thus, more appropriate methods for GS in polyploids could be evaluated, possibly improving trait prediction.

Polyploidy can affect phenotypes through allelic dosage (additive effect of multiple copies of the same alleles), or by creating more complex interactions between loci or alleles, such as dominance or epistasis (Osborn *et al.* 2003). Thus, the inclusion of allelic dosage information may improve GS results (e.g., better fit, increase of accuracy) by creating a more realistic representation of the effects of each genotypic class. Although the evidence of dosage effects in the expression of important economic traits exists (Guo *et al.* 1996; Birchler *et al.* 2001; Adams *et al.* 2003; Osborn *et al.* 2003), few studies linking dosage effects to phenotype prediction have been reported in autopolyploid species (e.g.; Slater *et al.*, 2016; Sverrisdottir el al. 2017; Nyine *et al.* 2018; Endelman *et al.* 2018). It is interesting to note that genotype classification is one of the major challenges for polyploids. Studies about genotyping calling evaluation for autopolyploids with next generation sequencing (NGS) data showed that none of the existing methods performs properly (Grandke *et al.* 2016), unless high sequencing coverage (60-80x) is used (Uitdewilligen *et al.* 2013).

Here we compare a novel approach to GS in the context of autopolyploid, using *Vaccinium corymbosum* (southern highbush blueberry, SHB) as a model. The cultivated SHB is an autotetraploid, presenting 2n = 4X = 48 chromosomes (Lyrene *et al.* 2002). Inbreeding depression is strong in SHB and population improvements have been achieved by long-term recurrent phenotypic selection alongside with long testing phase and slow genetic gain per generation (Lyrene 2008). Our goal was to investigate and compare the influence of different relationship matrices that consider different ploidy information on phenotype prediction, using novel genotyping approaches based on next-generation sequencing.

## MATERIAL AND METHODS

### Population and phenotyping

The population used in this study encompasses one cycle of the University of Florida blueberry breeding program’s recurrent selection, comprising 1,847 SHB individuals. This population was originated from 124 biparental controlled crosses, from 146 parents that presented superior phenotypic performance (cultivars and advanced stage of breeding). Phenotypic data of eight yield and fruit quality-related traits were collected during two production seasons (2014 and 2015), when the plants were 2.5 and 3.5 years of age. Yield (rated using a 1-5 scale), weight (g), firmness (g mm^−1^ of compression force), scar diameter (mm), fruit diameter (mm), flower bud density (reported as buds per 20 cm of shoot), soluble solids content (°Brix), and pH were evaluated. The last three traits were phenotyped only in one year – soluble solids content and pH were phenotyped in 2014 and flower buds in 2015.

Five berries (fully mature and presenting picking quality) were randomly sampled to compose the measurement of fruit traits for each individual. Fruit weight was measured using an analytical scale (CP2202S, Sartorious Corp., Bohemia, NY). The FirmTech II firmness tester (BioWorks Inc., Wamego, KS) was used to measure fruit diameter and firmness. The scar diameter was obtained by image analysis of the fruits using FIJI software (Schindelin *et al.* 2012). The number of flower buds was counted in the main cane upright shoot, in the top 20 cm. A digital pocket refractometer (Atago, U.S.A., Inc., Bellevue, WA) was used to obtain soluble solids measures from 300μl of berry juice. The pH was measured using a glass pH electrode (Mettler-Toldeo, Inc., Schwerzenbach, Switzerland). More details are provided by Amadeu *et al.* (2016), Cellon *et al.* (2018), and Ferrão *et al.* (2018).

### Genotyping

Genomic DNA was extracted and genotyped using sequence capture by Rapid Genomics (Gainesville, FL, USA). Polymorphisms were genotyped in genomic regions captured by 31,063 120-mer biotinylated probes, designed based on the 2013 blueberry draft genome sequence (Bian *et al.* 2014; Gupta *et al.* 2015). Sequencing was performed in the Illumina HiSeq2000 platform using 100 cycle paired-end sequencing. After trimming (quality score of 20), demultiplexing, and removing barcodes, reads were aligned to the draft genome using Mosaik v.2.2.3 (Lee *et al.* 2014). Genotypes were called using FreeBayes v.1.0.1 (Garrison and Marth 2012) considering the diploid and tetraploid options. Single-nucleotide polymorphisms (SNPs) were filtered considering i) minimum sequencing depth of 40 (average depth for the population); ii) minimum SNP quality score (QUAL) of 10; iii) only biallelic markers; iv) maximum population missing data of 0.5; and v) minor population allele frequency of 0.05. After filtering a total of 85,973 SNP were used in the GS analysis. Further information regarding population composition and genotyping approach were described in Ferrão *et al.* (2018). The genotypes for the diploid calling were coded as *0* (*AA*), *1* (*AB*), or *2* (*BB*). For the tetraploid parameterization they were coded as *0* (*AAAA*), *1* (*AAAB*), *2* (*AABB*), *3* (*ABBB*), and *4* (*BBBB*). A third parameterization (assumption-free method) was used, which considered allele ratio #*A*/(#*A* + #*a*), where #*A* is the allele count (sequencing depth) of the alternative allele and #*a* is the allele count of the reference allele. No dosage calling was performed in this model (File S1); these data varied continuously between *0* and *1*.

### Population genetics analysis

In order to compare the information captured by each genomic-based relationship matrix, we performed linkage disequilibrium (LD), and principal components (PC) analyses. Pearson correlation tests (r^2^) were performed for pairwise LD estimation among SNPs within scaffolds, considering draft reference genomes (Bian *et al.* 2014; Gupta et al, 2015). One SNP was randomly sampled per probe interval, and a total of 22,914 SNP were used in the analysis. LD was obtained for all marker-based scenarios: i) diploid (*G2*); ii) tetraploid (*G4*) and iii) ratio (*i.e.*, continuous genotypes; *Gr*). The LD decay over physical distance was determined as the mean distance at the LD threshold of r^2^ = 0.2. To compare the LD among scenarios, the mean distances (Kb) and their interval confidences at r^2^ = 0.2 were compared. The diversity captured from each relationship matrix was compared by PC using the R package adegenet v. 1.3-1 (Jombart and Ahmed 2011).

In order to compare the information present in the marker matrices, we also evaluated the observed heterozygosity in the population. For this, we obtained the ratio between the number of heterozygote genotypes and the total number of individuals. To estimate the heterozygosity for the continuous genotypes, empirical limits were established based on the mean and standard deviations presented for homozygotes classes of the tetraploid parameterization.

### Models

One-step single-trait Bayesian linear mixed models were used to predict breeding values for each individual in the population, as follows:

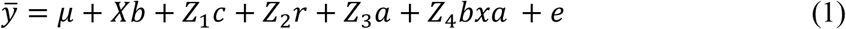

Where 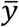 is a vector of the phenotypic values of the trait being analyzed, *μ* is the population’s overall mean, *b* is the fixed effect of year, *c* is the random effect of *i*th column position in the field 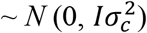, *r* is the random effect of the *i*th row position in the field ~ *N* 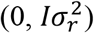, *a* is the random effect of genotype 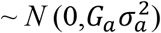, where *G*_*a*_ was replaced by the different additive relationship matrices as described in the next section. The *bxa* is the random effect of the year by genotype interaction 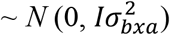, and *e* is the random residual effect 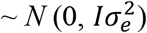. Row and column effects were considered nested within year only for the traits evaluated in two years. For traits measured a single year, the same equation (1) was used without the year and the year by genotype interactions. The variance components for each random variable were: additive 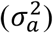, column 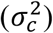, row 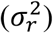, year-by-genotype interaction 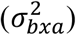, and residual 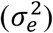. *X*, *Z*_1_, *Z*_2_, *Z*_3_, and *Z*_4_ were incidence matrices for year, column, row, genotype, and year by genotype interaction, respectively. The narrow-sense heritabilities were estimated considering the ratio between the additive variance component and the total phenotypic variance (sum of all variance components).

### Relationship matrices

To quantify the effect of the genetic information used to build the relationship matrices on the predictive ability (PA), we performed analyses considering different approaches to modeling the genotypic values in autotetraploid species (Table 1, File S1). The factors tested were: i) the source of information used to build the relationship matrix (pedigree, genomic, or no relationship information); and ii) ploidy information (diploid, tetraploid, and assumption-free method).

**Table 1.**
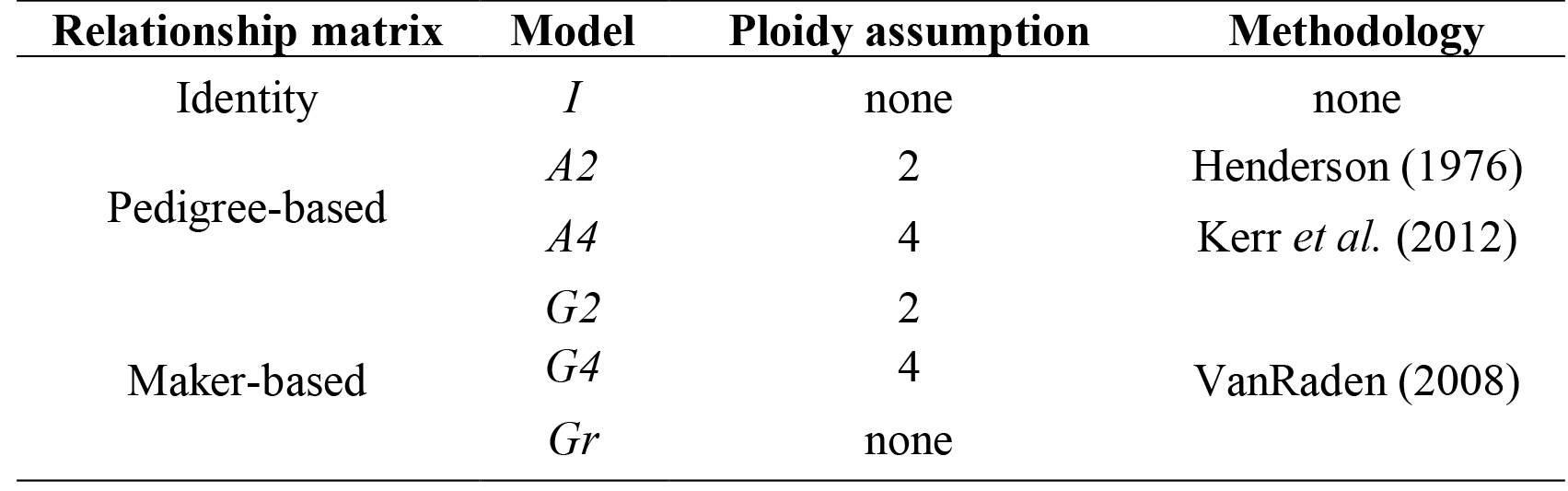
Methods and assumptions used to compare the influence of relationship matrices, ploidy and continuous genotypes in the prediction of breeding values for blueberry.

The methods chosen to obtain the relationship matrices are shown in the Table 1. The pedigree-based relationship matrices (*A*) were built considering a diploid model (Henderson 1976) and autotetraploid model without double-reduction (Kerr *et al.* 2012). The marker-based relationship matrices (*G*) were based on the incidence matrices of markers effects (*X*) according to VanRaden (2008) and adapted by Ashraf *et al.* (2016). Different assumptions can be made regarding the marker allele dosage in autotetraploids (Table 2). We built the *X* matrices under three assumptions regarding the additive marker allele dosage effect: *i)* a pseudo-diploid model, where all the heterozygous genotypes are assumed as one class, corresponding to a unique effect (data coded as *0*, *1*, and *2*); *ii)* an additive autotetraploid model, where each genotype had a specific value, and cumulative additive effect is assumed (data coded as *0*, *1*, *2*, *3*, and *4*); and *iii)* an assumption-free method based on the ratio of reads count for the alternative and reference alleles (continuous parameterization, assuming values between *0* and *1*), where also a cumulative additive effect is assumed. For the construction of the relationship matrices based on marker data, the missing genotypes were substituted by the mean. The R package AGHmatrix v. 0.0.3003 (Amadeu *et al.* 2016) was used to obtain all relationship matrices.

**Table 2.** Genotype codes for marker-allele dosage effects with different assumptions. Adapted from Slater *et al.* (2016).

| Genotype | Pseudo-Diploid | Autotetraploid | Continuous values* |
| --- | --- | --- | --- |
| AAAA | 0 | 0 |  |
| AAAB | 1 | 1 |  |
| AABB | 1 | 2 | 0 - 1 |
| ABBB | 1 | 3 |  |
| BBBB | 2 | 4 |  |

\* Continuous values with a ploidy assumption-free parameterization

### Model implementation

The six models described above (Table 1) were fitted using the R package (R Core Team 2018) BGLR v. 1.0.5. (de los Campos and Pérez-Rodríguez 2016). The predictions were based on 30,000 iterations of the Gibbs sampler, in which 5,000 were taken as burn-in, and a thinning of five. The number of iterations, burn-in, and thinning interval parameters were evaluated to define the final values used in the analysis (Figure S1). A single step regression approach was applied to perform all phenotypic BLUP (I matrix), pedigree-BLUP (P-BLUP), and genomic-BLUP (G-BLUP). Default hyper-parameters were previously described (Perez and de los Campos 2014).

### Validation and model comparison

For each trait, models were compared based on their PA, stability (mean square errors), goodness-of-fit, and expected genetic gain. A 10-fold cross validation scheme was applied to compute model PA. Because each validation group might have a different mean (Resende *et al.* 2012b), the phenotypic PA were obtained as the Pearson correlation coefficient between the empirical best linear unbiased estimation values (eBLUEs) obtained by considering all the variables in the equations 1 as fixed (*i.e.*, Least Square means estimations; LSMeans) and the cross-validated breeding values (BV) predicted by the models for each validation fold. The goodness-of-fit for the different models was evaluated with measures of the posterior mean of the log likelihood obtained from the full data set. The model with the lowest value for this parameter defined the best fit for the data. For the expected genetic gain estimation we used the following formula: ΔG=(*PA* · *σ*_*a*_ · *i*)/*L*, where *PA* is the phenotypic predictive ability, *σ*_*a*_ is the square root of additive genetic variance in the population, *i* is the selection intensity, and *L* is the breeding cycle length. The selection intensity (*i*) was considered constant for all methods.

Phenotypic and genotypic data used for diploid and tetraploid parameterizations are available from Dyrad Digital Repository (accession number doi: 10.5061/dryad.kd4jq6h). Data for continuous parameterization is available for review upon request. Data will be available at Dyrad Digital Repository. The authors affirm that all data necessary for confirming the conclusions of the article are present within the article, figures, and tables.

## RESULTS

### Population genetics analysis

Linkage disequilibrium decayed below r^2^ = 0.2 at distances of 88.3 Kb, 92.6 Kb, and 98.2 Kb for the diploid, tetraploid and continuous models, respectively (Figure 1A-C). No significant difference was observed considering the confidence interval for the mean distance (Kb) at r^2^ = 0.2 among different ploidies and continuous genotyping scenarios (Figure S2).

**Figure 1.**
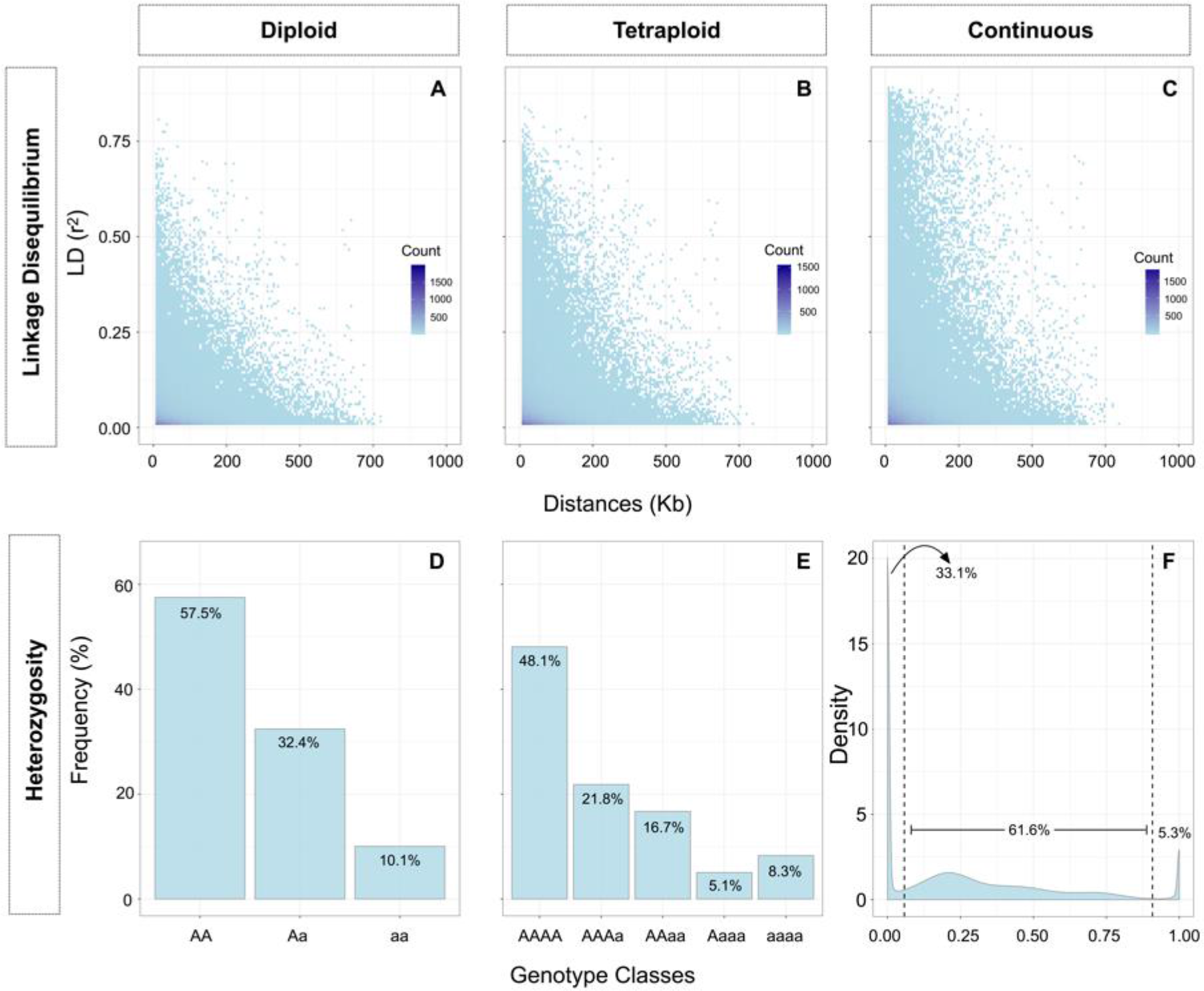
Linkage disequilibrium decay and heterozygocity for blueberry. Linkage disequilibrium decay estimation using one marker per probe, within scaffolds for (**A**) diploid, (**B**) tetraploid and (**C**) continuous genotype parameterizations. Heterozygosity observed in (**D**) diploid, (**E**) tetraploid, and (**F**) heterozygosity empirically established for the continuous genotypes scenario, assuming the limits of 0.058 ≤ X ≤ 0.908.

Similarly, no major differences were found between parameterizations within methodology (*i.e.*, pedigree-based or marker-based methods) in the PC analysis (Figure S3). The first two PC components of the marker-based (*G*) matrices were consistent across all matrices, explaining approximately 20% of the variation. For example, *G2* matrix captured 20.60% of the variation, while *G4* captured 21.71%, and *Gr* captured 23.36% (Figure S3 A-C). The PC results were consistent between pedigree methodologies as well. Approximately 38% of the variation was explained (*i.e.*, 37.74% of the variability was explained for the *A2* matrix and 37.86% was explained for the *A4* matrix, Figure S3 D-E). The results obtained in the PC analysis did not justify a stratified sampling of cross-validation populations, since no evidence of sub-population structure was detected for any of the relationship matrices.

Considering the heterozygosity observed in each scenario, genotypes assumed as homozygotes in the diploid parameterization were classified as one of the possible heterozygote classes in the tetraploid and in the assumption-free parameterizations (Figure 1D-F). As a result of this process, the tetraploid parameterization presented 37.50% more heterozygotes than the diploid parameterization. Considering the empirical thresholds established to compare the proportion of “heterozygotes” in the continuous genotypes with the ploidy parameterizations, values equal to or below 0.058 and equal to or above 0.908 were considered as “homozygotes” classes (dashed lines, Figure 1F). With this, 61.59% of the genotypes were considered “heterozygotes”, thus the continuous method would have presented 89.92% and 41.23% more heterozygotes than the diploid and the tetraploid parameterization, respectively. Nevertheless, some misclassification of data into classes in the diploid and tetraploid parameterization might have occurred (Figure 2A-B).

**Figure 2.**
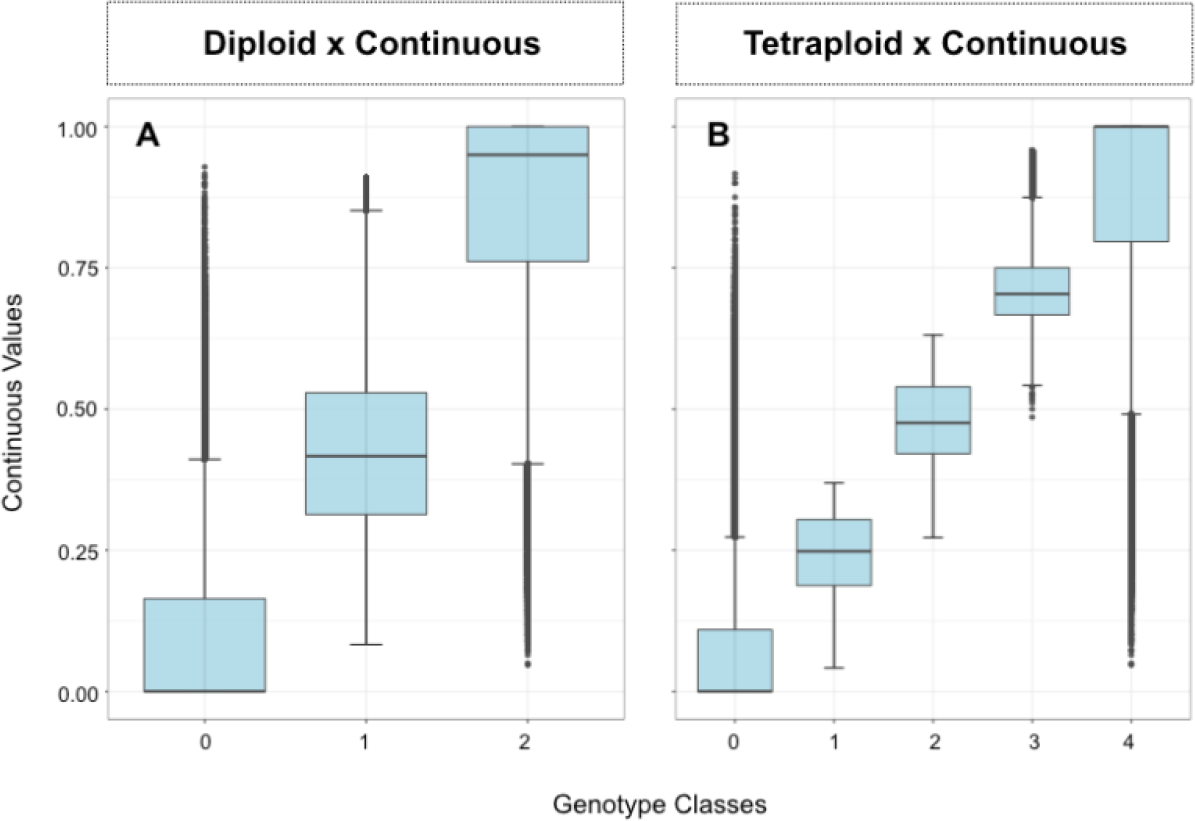
Relationship of the continuous values considering the classes assumed in the (**A**) diploid and (**B**) tetraploid parameterizations.

### Variance estimates

The posterior means of the genetic parameters are summarized in Table 3. All the traits presented additive genetic variance significantly higher than zero. A wide range of variance was observed within a given parameter for the different methodologies, and most of the values were significantly different from each other (considering Tukey test results; Table 3, Table S1). Marker-based methodologies generated significantly smaller estimations for variance components when compared with pedigree-based estimations. Within marker-based methodologies, the assumption-free parameterization generated significantly smaller estimations. The effects of the difference in the estimation of variance components are reflected in the estimated heritabilities – smaller values were estimated for marker-based methodologies. The lowest heritability was obtained for soluble solids, flower buds, and pH. Considering all methods, narrow-sense heritability values varied between 0.152 and 0.574, for flower buds and fruit weight, respectively.

**Table 3.**
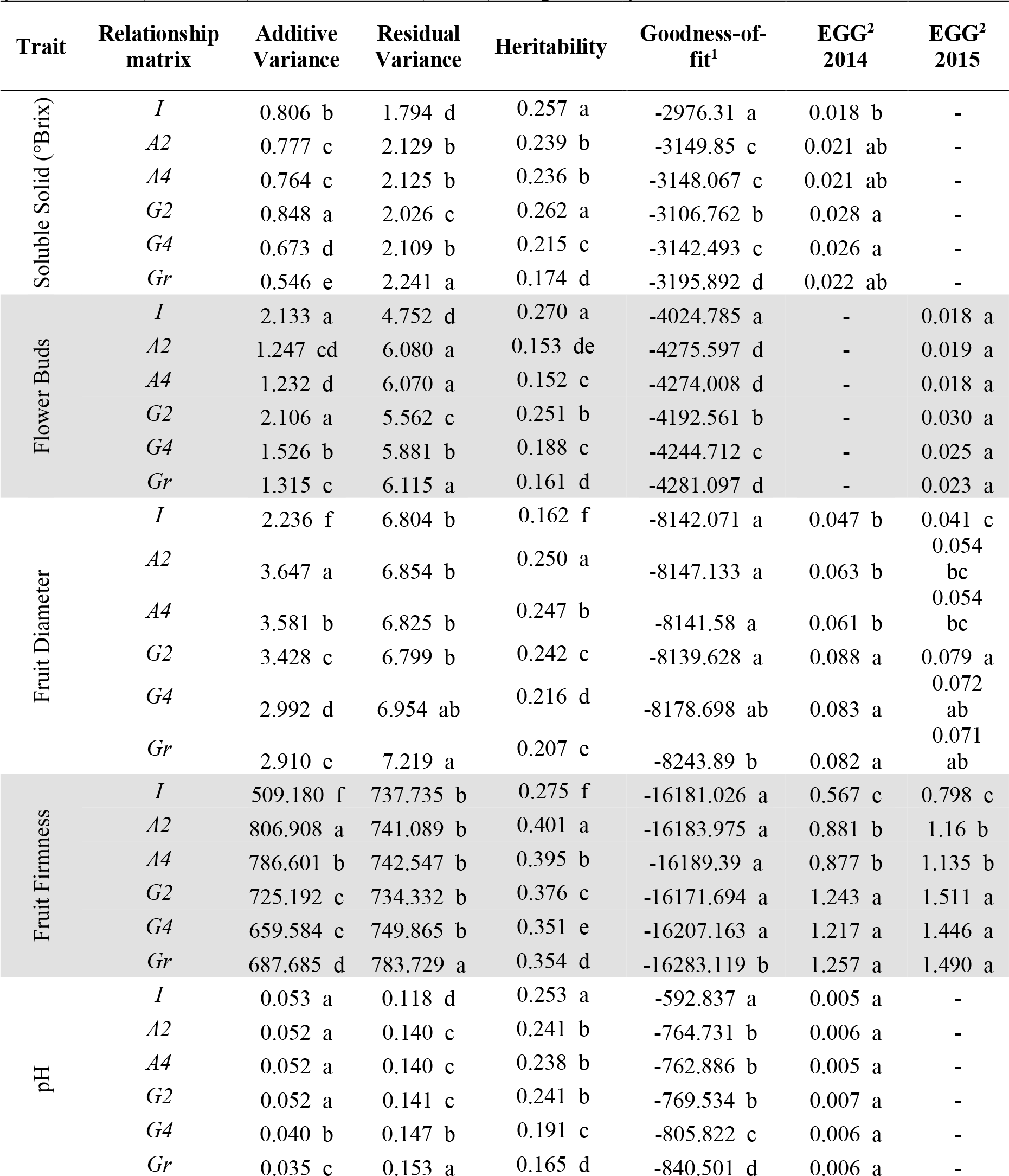
Genetic parameters estimated for eight yield and fruit-related traits analyzed with six linear mixed models, considering the use of ploidy information and continuous genotypes. Source of information, and dosage parameterizations for the relationship matrices indicated by the letters (*I*, *A*, or *G*), and numbers (2 or 4), respectively*

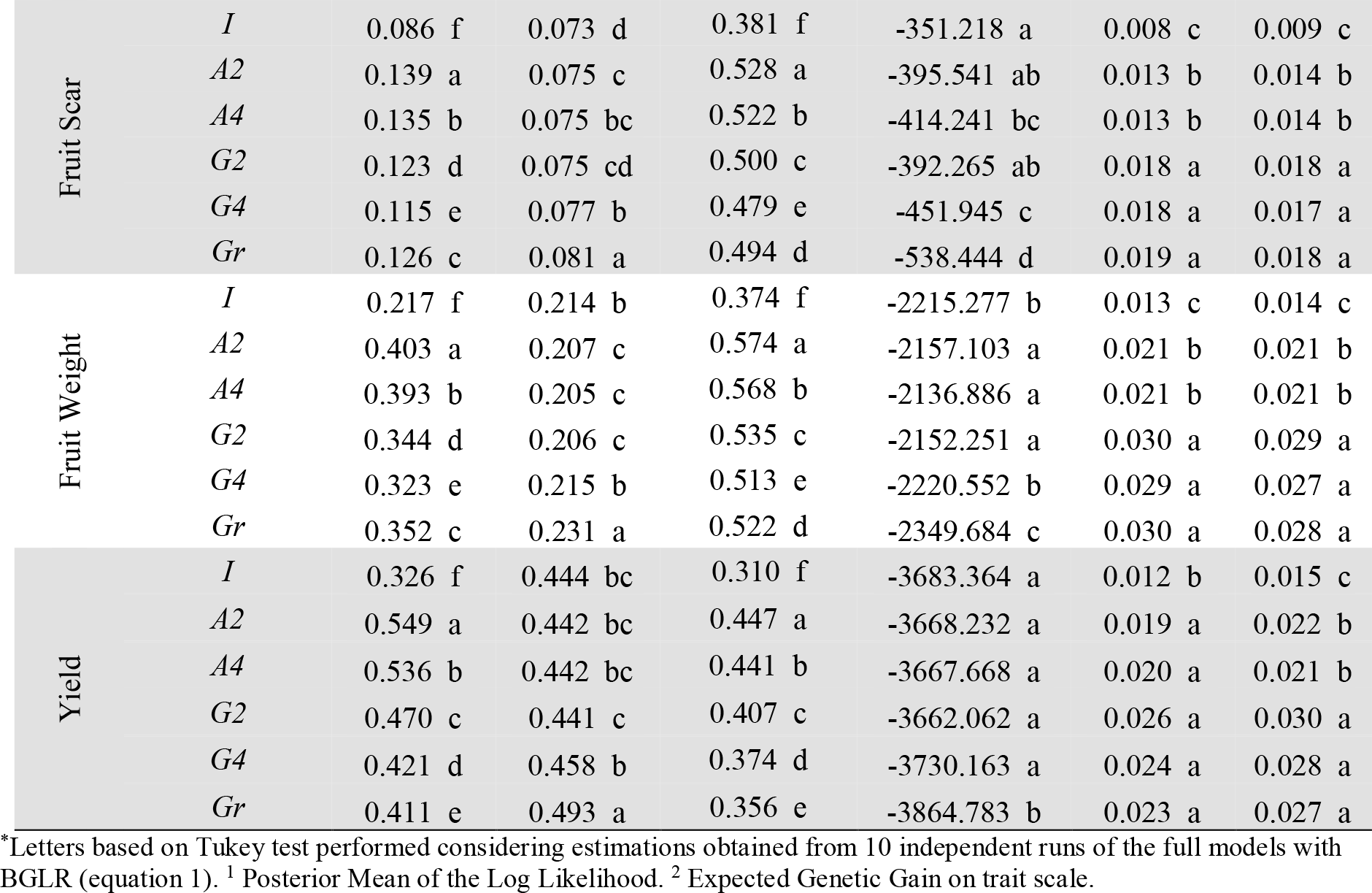

### Effect of the genetic information to build the relationship matrices

The incorporation of relationship information in the analysis generated better PA results than the phenotypic-BLUP model without it. Overall, we observed that higher values for the phenotypic PA were obtained when marker-based relationship matrices were used, when compared with phenotypic and pedigree BLUP (*I* and *A* matrices, respectively). However, the marker-based and pedigree-based results were not always significantly different from each other (Figure 3, Table S1). The use of molecular data yielded phenotypic PA values ranging from 0.27 (pH) to 0.49 (fruit scar) in 2014, and from 0.15 (flower buds) to 0.51 (fruit firmness) in 2015. Lower PA values were obtained for traits with lower heritability and better results were observed for the second year of evaluation. The biggest increase in the PA values can be seen for fruit firmness – when we compared marker and pedigree results, we observed an average increase of 13.37% in 2014. Also, an increase in the PA values of 11% was observed for fruit diameter and yield in 2015 when markers were used instead of pedigree data.

**Figure 3.**
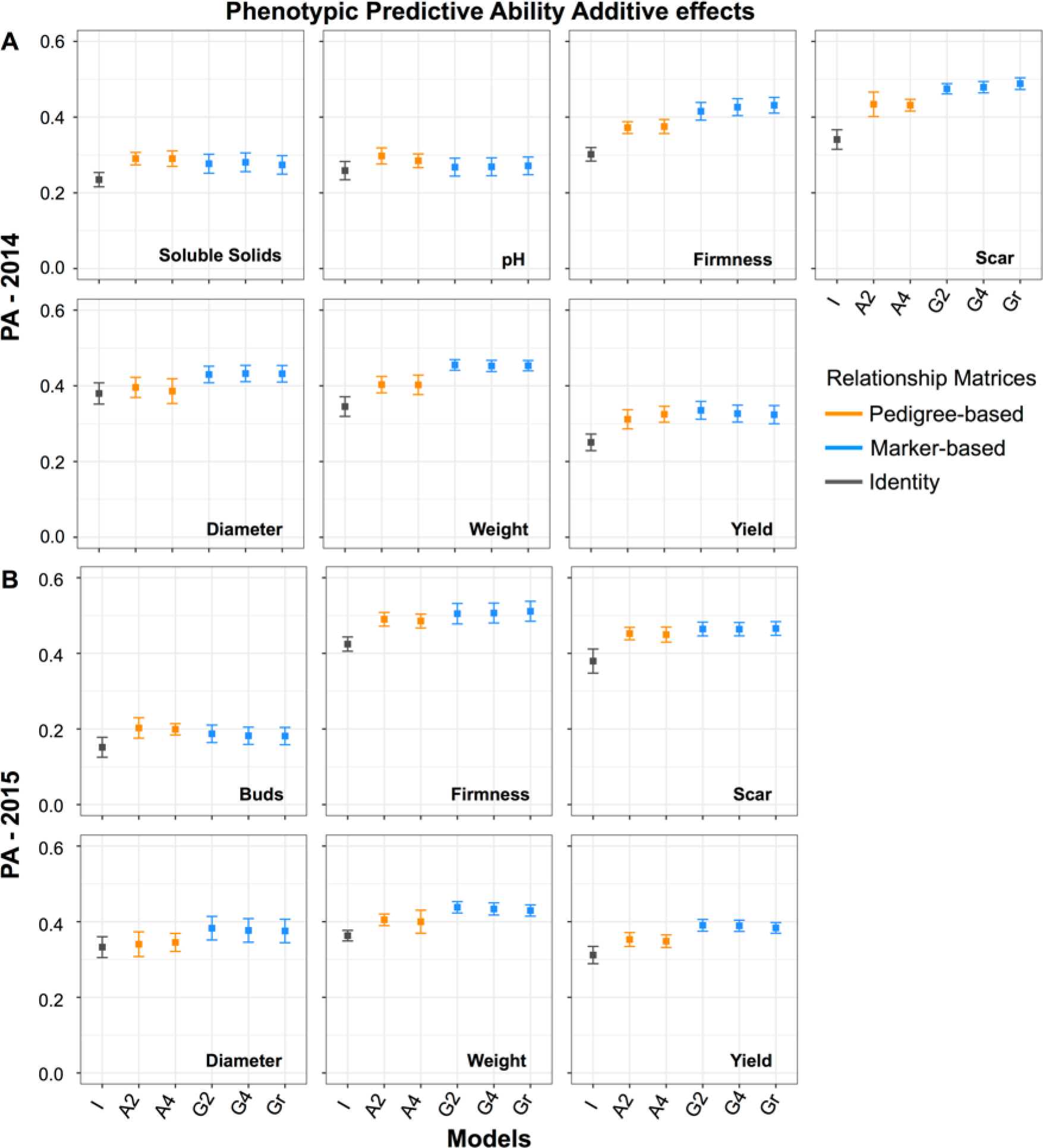
Phenotypic predictive abilities for (**A**) seven traits in 2014, and (**B**) for six traits in 2015 for different dosage parameterizations (indicated by the numbers 2 or 4) of the relationship matrices (indicated by the letters *I*, *A*, and *G*) in the prediction of breeding values of 1,847 SHB individuals.

The use of pedigree-based relationship matrices generated higher phenotypic PA values for all the traits, when compared with the assumption of unrelated individuals (*i.e.*, identity matrix). Unlike the identity matrix, the use of pedigree-based matrix assumes that there is relationship (expected values) among individuals. The phenotypic PA obtained for the pedigree methods in 2014 yielded values from 0.20 (flower bud) to 0.49 (fruit firmness). As with marker-based methods, smaller values were observed for traits with lower heritability (*i.e.*, pH, brix, and flower bud). For 2015, the PA results for the phenotypic-BLUP were 0.36, 0.38, and 0.42, for fruit weight, fruit scar, and fruit firmness, respectively. The PA values obtained for the same traits with pedigree-BLUP were 0.40, 0.45, and 0.49, respectively. No significant differences between the models’ stability were observed (Table S1).

### Use of dosage information and continuous genotypes

Our results indicate that the importance of dosage in GS will vary depending on the trait being analyzed. For example, in 2014 the PA for fruit firmness, fruit scar, and fruit diameter showed modestly better phenotypic PA when the tetraploid and continuous parameterizations were applied, as opposed to the diploid parameterization (Figure 3, Table S1). Although no significant difference was observed between marker-based models, the use of relationship matrices derived from continuous genotype data (ploidy-free parameterization) performed equally well as the best models (Figure 3, Table S1). However, the goodness-of-fit statistics show that the use of a relationship matrix obtained from the continuous genotype data significantly improved model fit for all traits (Table 3). This was followed by the tetraploid parameterization using marker-based data.

### Expected genetic gain in a perennial fruit tree, blueberry

The results obtained for the expected genetic gain (EGG) are summarized in Table 3. GS offers the possibility to accelerate genetic improvement by decreasing the breeding cycle and selecting superior individuals earlier in the breeding program. Considering a breeding cycle (*L*) of 12 years (Cellon *et al.* 2018) we propose that routine genomic selection could be implemented in the second stage of the blueberry breeding program, which would allow the omission of a whole stage (stage III), and a three-year reduction for cultivar release (Figure 4).

**Figure 4.**
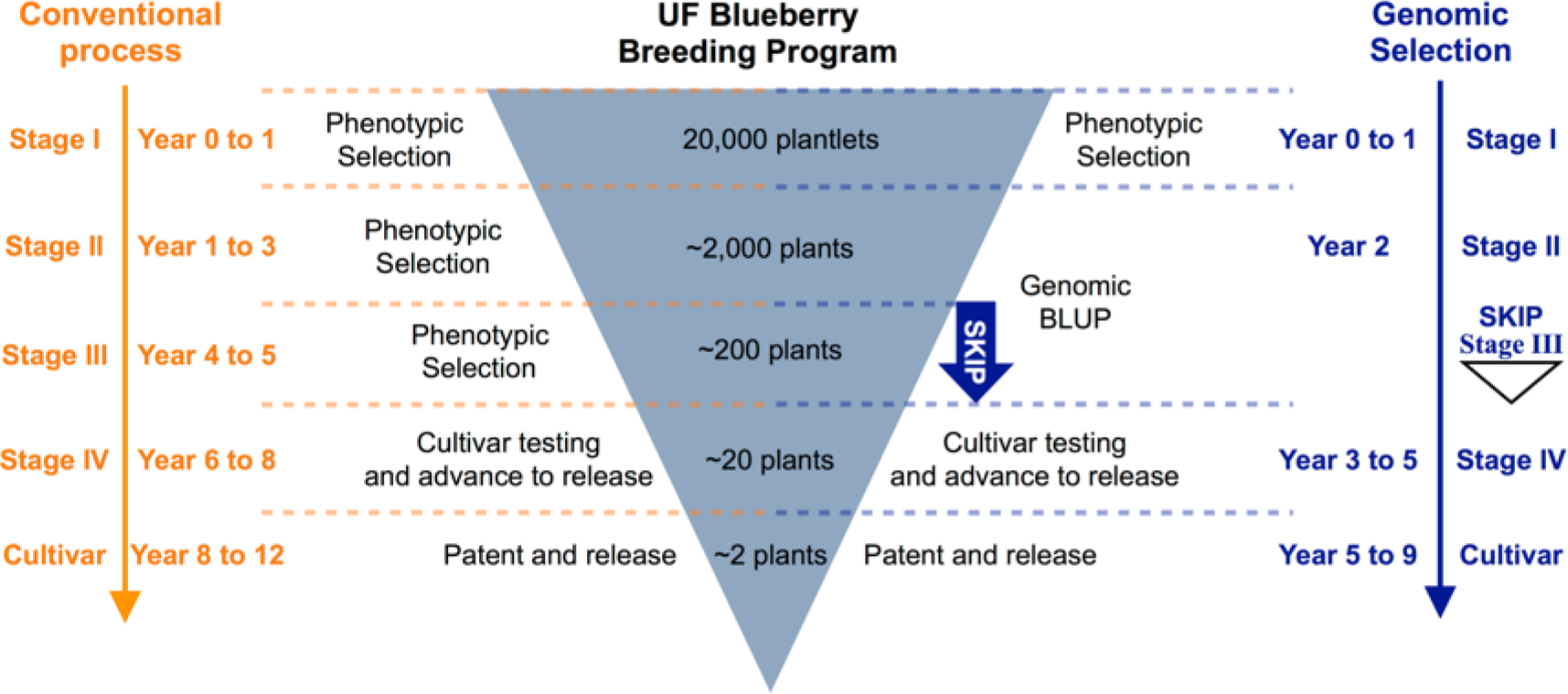
Example of the University of Florida blueberry breeding program stages and times of selection. Conventional process (left) compared with the proposed process implementing genomic selection (right).

Higher EGG was obtained for all traits when marker-based matrices (*i.e.*, genomic selection) were applied (Table 3), which was mainly related to the reduction in cycle time. The implementation of GS in the second stage population would lead to an increase in the EGG varying from 27% (pH) to 119% (scar) when compared with the application of phenotypic BLUP. Considering the comparison of marker-based and pedigree-based models, an increase of 15% (pH) to 41% (fruit weight, fruit scar, and flower buds) in the EGG was observed (Table 3). In addition, the use of continuous data generated EGG values that were not significantly different of the best models for all traits (Table 3).

## DISCUSSION

In this study, six linear mixed models were applied to predict breeding values for eight yield and fruit-quality traits measured in a real blueberry breeding population as model. Analyses were based on phenotypic, pedigree, and high-density marker data from 1,847 individuals. We compared the expected genetic gain, the stability, and the PA of models considering different sources to build the relationship matrices (only phenotype=BLUP, phenotypes + pedigree=P-BLUP, phenotypes + genomic=G-BLUP). Our results also explored models accounting for ploidy information and proposed the use of genotypic data that is independent of assumptions regarding ploidy levels (continuous) to perform GS, avoiding the need for *a priori* parameterization for a given ploidy level.

### Continuous data

Our research showed empirical evidences that the use of continuous genotypic data from NGS can be effectively applied in GS models for autotetraploid species. This method was tested and compared with marker calling methodologies at the individual level in genome wide association studies (Grandke *et al.* 2016). It was also tested in family pool data for GS (Ashraf *et al.* 2014; Guo *et al.* 2018), as well as used at the individual level in tetraploid potato for GS by Sverrisdottir *el al.* (2017). However, to our knowledge the comparison of continuous genotypes with ploidy parameterizations for genomic selection has not yet been reported. Here we empirically compare diploid, tetraploid, and continuous data at the individual level for the application of genomic selection in an autotetraploid species.

In polyploids, the assignment of genotypic classes based on NGS data is a major challenge, with high risk of misclassification (Grandke *et al.* 2016, Bourke *et al.* 2018). The problem is further exacerbated as the ploidy increases – for a given level of ploidy, *n*, the expected number of genotypic classes is *2n*+*1*. As a consequence, the signal distribution derived from each genotypic class increasingly approximates a continuous distribution where no clear separation is observed (Grandke *et al.* 2016). Despite extensive research to address these challenges (Serang *et al.* 2012), advances have been mostly limited to SNP arrays in tetraploid data (Carley *et al.* 2017). Studies that evaluated genotype calling with NGS data obtained from polyploids show that no method works properly, and that misclassification of genotypes can significantly interfere in the results of genetic studies (Grandke *et al.* 2016). This misclassification can be observed in our results when a diploid, or tetraploid parameterization is used in the genomic data (Figure 2A-B) with standard parameters of filtering. The use of the continuous genotyping approach provides a relevant alternative to overcome this issue that is independent of assumptions regarding ploidy level. Models that used continuous genotypic data performed as well as the best models and resulted in modestly better predictive abilities for some of the traits (*i.e.*, fruit firmness, fruit scar, and fruit diameter; Table 3), but better data fit, which could indicate better prediction of future populations. The use of continuous genotypes also simplifies the analysis complexity and time by eliminating the genotype calling and parameterization for a give ploidy, because instead, the ratio of reads assigned to each allele are used. Finally, our results showed that the addition of noise associated with the continuous distribution in the genotypes significantly improved model fitting for all analyzed traits (Table 3), instead of increasing the complexity of the models. The benefits of continuous genotyping could easily be extended to more complex polyploids (higher ploidies), where the genotype attribution is even more difficult, however higher sequencing depth would probably be required. Meanwhile, for more complex models, such as those that consider dominance effects, dosage calling is still necessary.

### Relationship matrices

Our results also showed that including information based on the genetic merit of the individuals yielded better results when compared with the phenotypic-BLUP analysis (based on the identity matrix; Table 3), corroborating previous studies in the literature (e.g., Muir 2005; Resende *et al.* 2012a; Muñoz *et al.* 2014a). In addition, the use of marker-based methodologies generated better predictions than pedigree for most of the traits. Marker-based methods allow the capture of Mendelian segregation. This is especially important in our population, since it was composed of 117 full-sib families. In this context, pedigree-based methods have no power to distinguish variance within families. Another advantage is that marker-based methods allows the computation of genetic similarity among unidentified individuals in the pedigree, and corrections of errors in the pedigree, which can affect parameter estimation causing reduction in the genetic gain (Muñoz *et al.* 2014b).

In our results, some non-significant differences between pedigree and marker-based methods were identified, which could be an effect of the extensive pedigree data used, as well as bias in pedigree-based estimations. Pedigree-based methods can overestimate the reliability of selection and consequently, the accuracy (Bulmer 1971; Gorjanc *et al.* 2015). Furthermore, it also presents low efficiency to capture and estimate genetic relationships among individuals (Resende *et al.* 2017).

It is interesting to notice that we used extensive pedigree information that dates back to 1907 for our predictions, which may not be common in other autopolyploid breeding. This extensive information can have significant implications on the estimation of relationship coefficients (Amadeu *et al.* 2016) and consequently, in breeding value predictions. For breeding programs with smaller pedigree depth information, the comparison between accuracies of prediction from marker and pedigree-based methodologies could be even bigger than what was found in our study.

### Allele dosage

The results obtained for both models that assumed more than three genotypic classes (*G4* and *Gr*) demonstrate the importance of considering dosage in the prediction of breeding values. However, this will depend on the trait analyzed, as previously reported by Nyine *et al.* (2018) and Endelman *et al.* (2018). For example, modest improvement was verified in the PA for fruit firmness, fruit scar, and fruit diameter when this factor was considered in the models.

In addition, model fitting was significantly better for methods that accounted for dosage information (Figure 3, Table 3, Table S1). The inclusion of nonadditive effects into the models could also improve model accuracy. Endelman *et al.* (2018) demonstrated that the inclusion of digenic effects, as well as accounting for ploidy information, presented a higher accuracy over diploid models when using a SNP array.

### Genomic selection for perennial autopolyploids

We also demonstrate the value of applying GS in a perennial fruit tree, blueberry. One cycle of blueberry breeding takes from 12 to 15 years until the release of a new cultivar (Lyrene 2008; Cellon *et al.* 2018). By applying selection based on high-density markers at early stages of the program, the time to cultivar release could decrease by three years (Figure 4), significantly improving the expected genetic gain per unit of time. More specifically, the use of GS would lead to an average increase of 86% in the EGG when compared with phenotypic BLUP, and an average increase of 32% over the application of pedigree-based models (Table 3). Implementing GS in this form could eliminate one stage in the breeding and selection process toward cultivar development, which will reduce costs associated with field trials and phenotyping. The implementation of GS would require extra financial outlay when genotyping and accurately phenotyping the training population. However, the savings on phenotyping and field trials of future generations (selection populations) could results in a break-even financial exercise, and as a result could be a cost-effective application of GS. However, this financial analysis needs to be performed for each crop in a case-by-case basis.

## Acknowledgments

The authors thank the University of Florida blueberry breeding program technical support, especially Dr. Paul M. Lyrene, David Norden, and Werner Collante. Special thanks to James Olmstead and Catherine Cellon, who coordinate the phenotyping and genotyping of the population as part of Catherine Cellon MS degree.

## Supplemental Material

**Figure S1.**
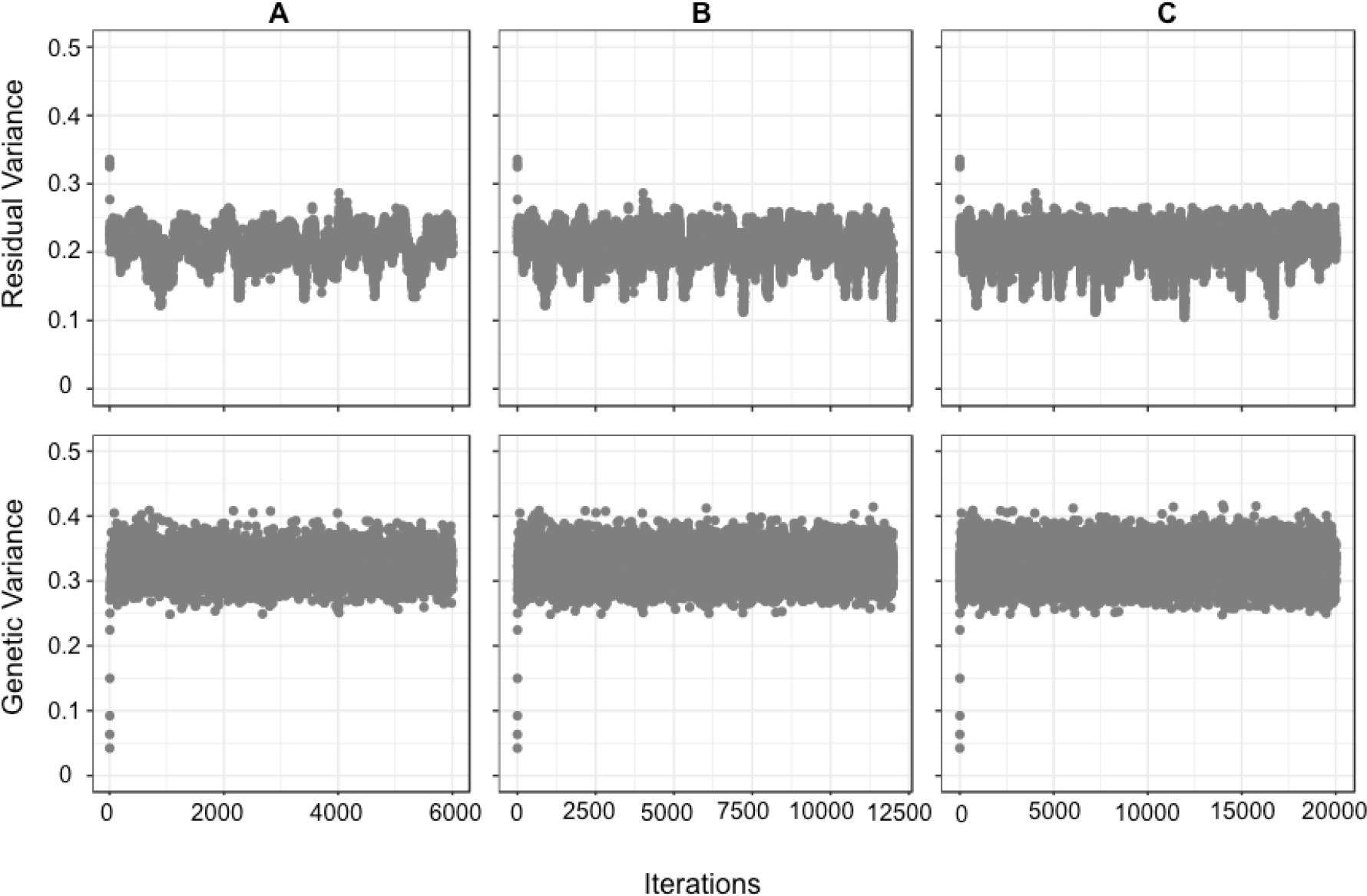
Model convergence obtained using different values for iterations (I), burn-in (B) and thinning (T) to perform the GS in BGLR for 1847 SHB genotypes. **A**) I=30K B=5K and T =5; **B**) I=60K B=10K and T =5; **C**) I=100K B=10K and T =5.

**Figure S2.**
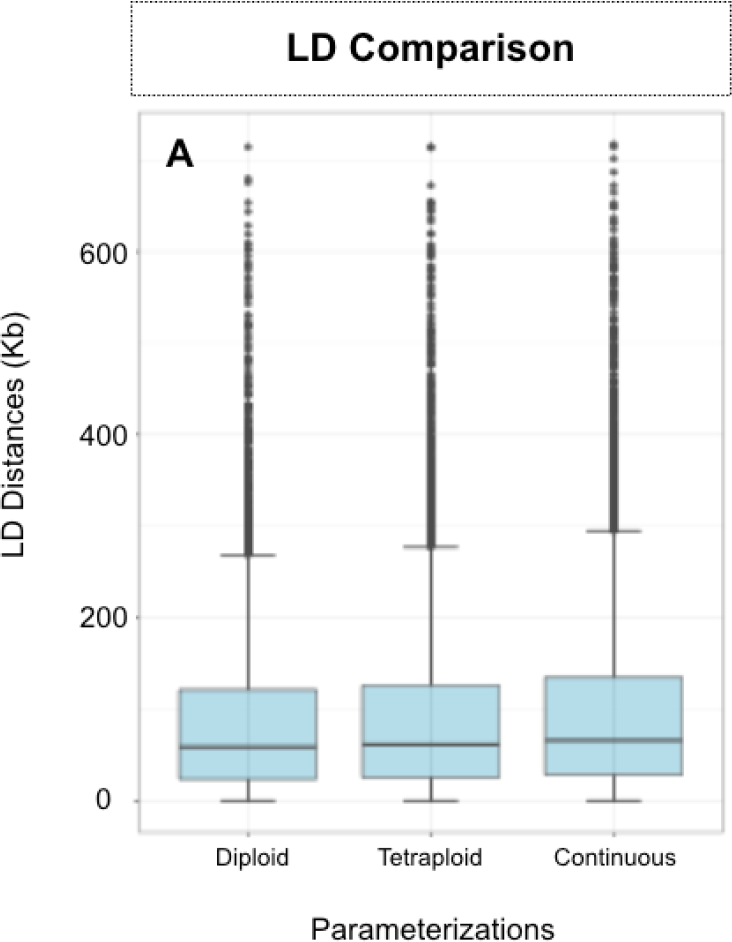
Distribution of linkage disequilibrium distances at the empirical threshold r^2^=0.2 for the diploid, tetraploid and continuous parameterizations.

**Figure S3.**
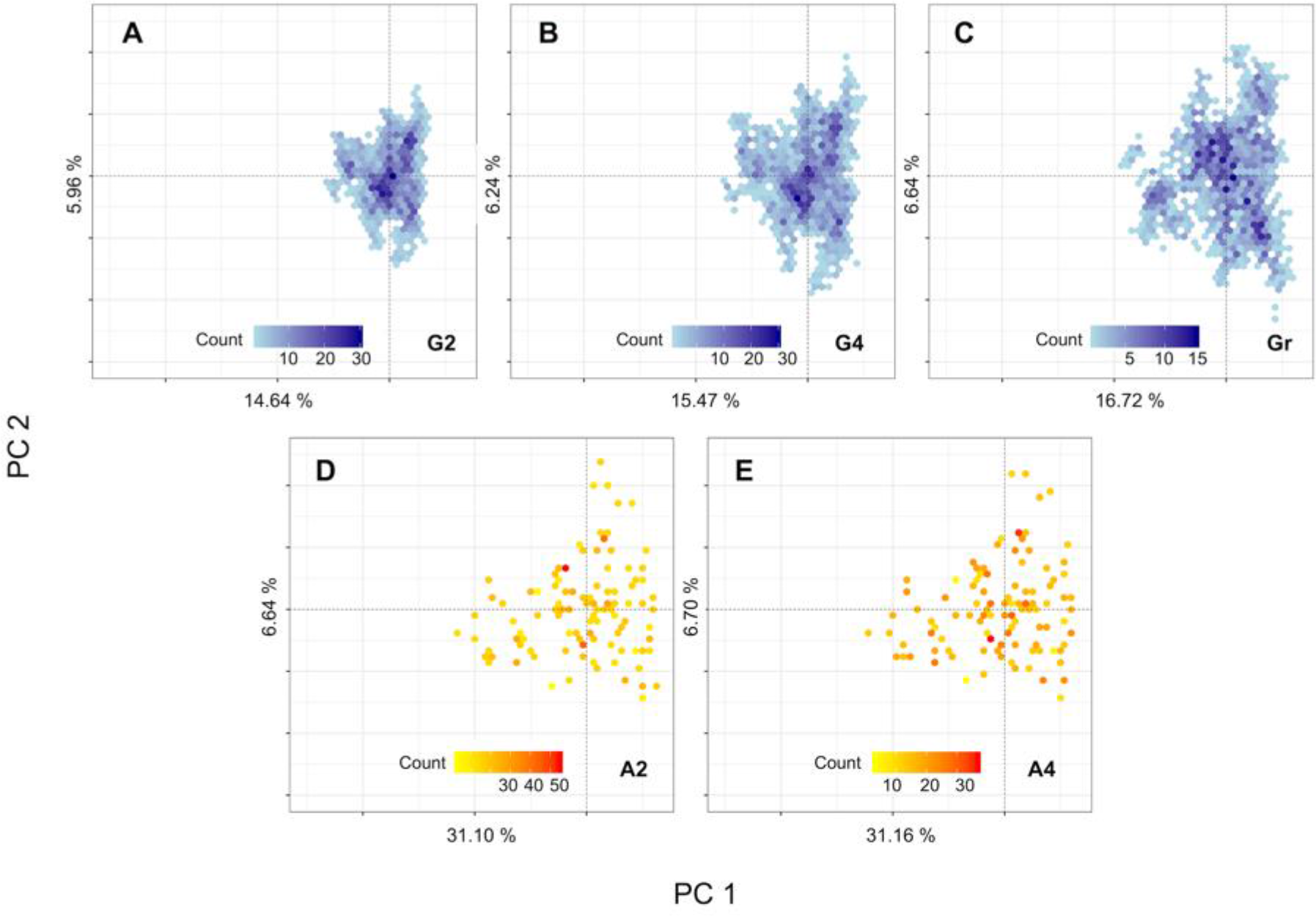
Principal components heat plots for the relationship matrices used in this study. In blue are the marker-based, and in orange are the pedigree-based matrices. Genomic matrices were computed using VanRaden’s (2008) methodology (**A**) Diploid, (**B**) tetraploid, and (**C**) continuous relationship matrix. Pedigree-based matrices were computed for the additive effects using (**D**) Henderson (1976) methodology for diploid, and (**E**) Kerr *et al.* (2012) methodology for tetraploid.

**Table S1.**
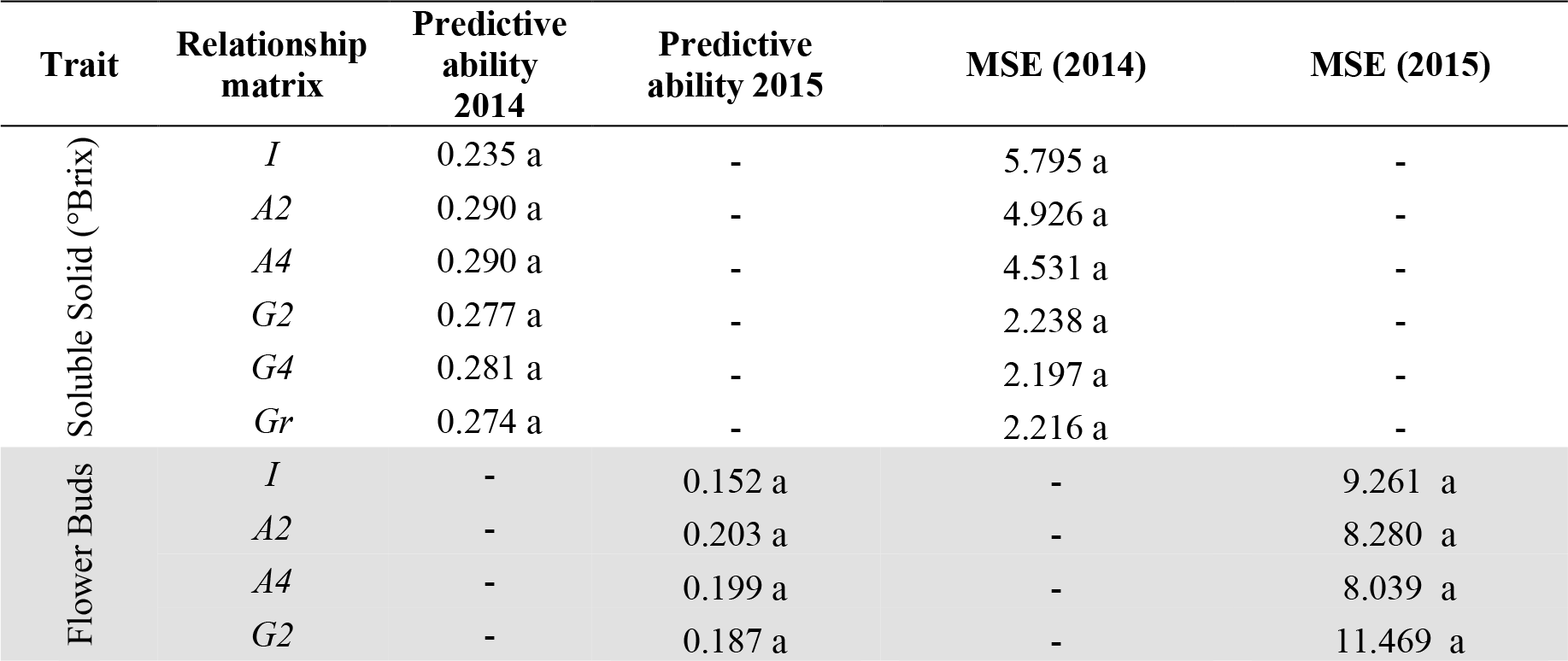
Accuracy and model stability values for eight yield and fruit-related traits analyzed with six linear mixed models with different dosage parameterizations of the relationship matrices. Source of information, and dosage parameterizations for the relationship matrices indicated by the letters (*I*, *A*, or *G*), and numbers (2 or 4), respectively*

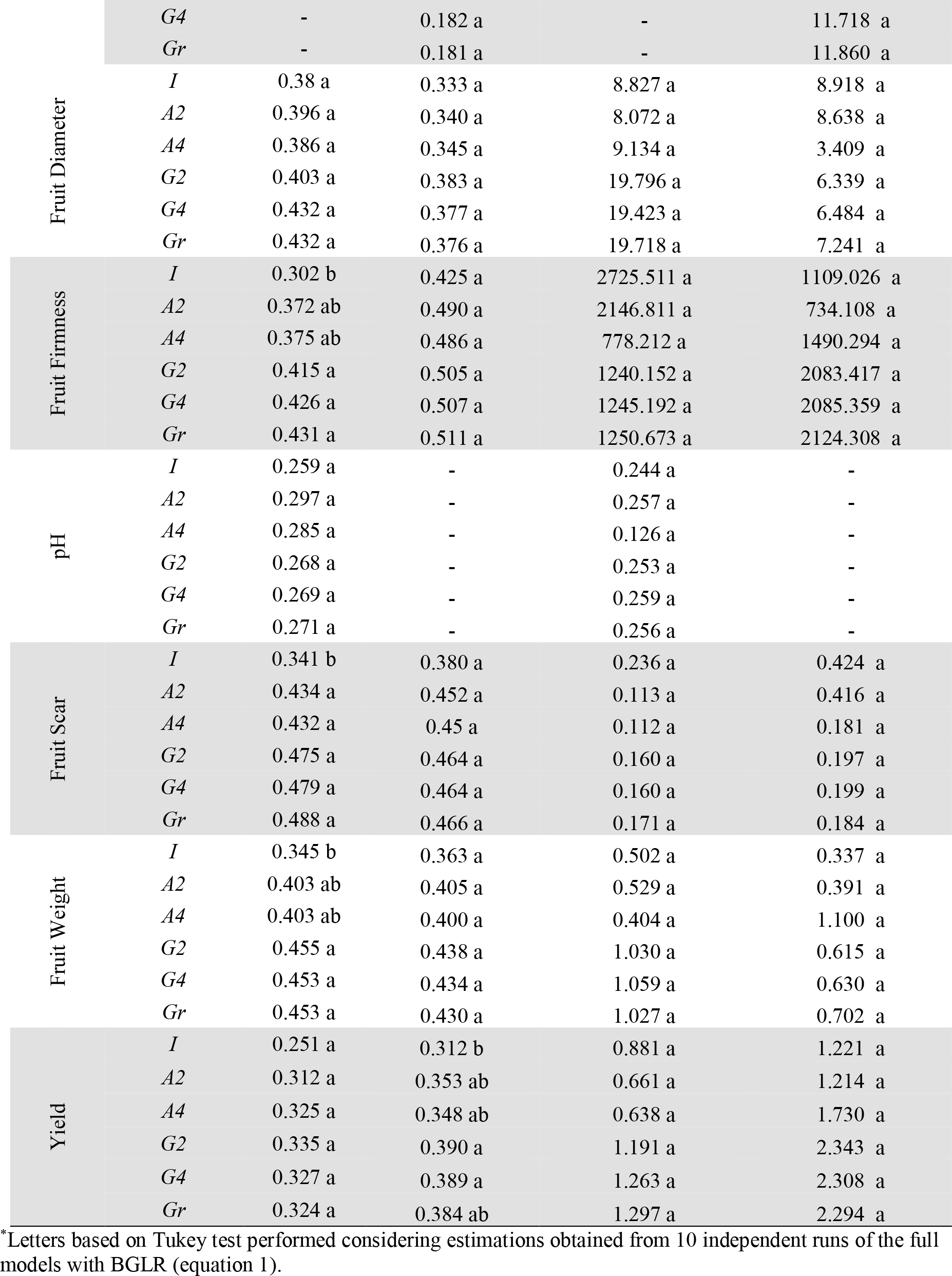

## Literature Cited

Adams, K. L., R. Cronn, R. Percifield, and J. F. Wendel, 2003 Genes duplicated by polyploidy show unequal contributions to the transcriptome and organ-specific reciprocal silencing. Proc. Natl. Acad. Sci. 100: 4649–4654.

Amadeu, R. R., C. Cellon, J. W. Olmstead, A. A. Garcia, M. F. Resende et al., 2016 AGHmatrix: R Package to construct relationship matrices for autotetraploid and diploid species: a blueberry example. Plant Genome 9: 3.

Annicchiarico, P., N. Nazzicari, X. Li, Y. Wei, L. Pecetti et al., 2015 Accuracy of genomic selection for alfalfa biomass yield in different reference populations. BMC genomics, 16: 1020.

Ashraf, B., V. Edriss, D. Akdemir, E. Autrique, D. Bonnett et al., 2016. Genomic prediction using phenotypes from pedigreed lines with no marker data. Crop Sci. 56: 957–964.

Ashraf, B. H., J. Jensen, T. Asp, and L. L. Janss, 2014 Association studies using family pools of outcrossing crops based on allele-frequency estimates from DNA sequencing. Theor. Appl. Genet. 127:1331–1341.

Bian, Y., J. Ballington, A. Raja, C. Brouwer, R. Reid, M. Burke et al., 2014 Patterns of simple sequence repeats in cultivated blueberries (Vaccinium section Cyanococcus spp.) and their use in revealing genetic diversity and population structure. Mol. Breed. 34: 675–689.

Birchler, J. A., U. Bhadra, M. P. Bhadra, and D. L. Auger, 2001 Dosage-dependent gene regulation in multicellular eukaryotes: implications for dosage compensation, aneuploid syndromes, and quantitative traits. Dev. Biol. 234: 275–288.

Bourke, P. M., R. E. Voorrips, R. G. Visser, and C. Maliepaard, 2018 Tools for genetic studies in experimental populations of polyploids. Front. Plant Sci. 9: 513.

Bulmer, M., 1971 The effect of selection on genetic variability. Am. Nat. 105: 201–211.

Carley, C. A. S., J. J. Coombs, D. S. Douches, P. C. Bethke, J. P. Palta et al., 2017. Automated tetraploid genotype calling by hierarchical clustering. Theor. Appl. Genet. 130:717–726.

Cellon, C., R. R. Amadeu, J. W. Olmstead, M. R. Mattia, L. F. V. Ferrao et al., 2018 Estimation of genetic parameters and prediction of breeding values in an autotetraploid blueberry breeding population with extensive pedigree data. Euphytica, 214: 1–13.

Comai, L., 2005 The advantages and disadvantages of being polyploid. Nat. Rev. Genet. 6: 836.

de Los Campos, G., and P. Perez-Rodriguez, 2016 BGLR: Bayesian Generalized Linear Regression. R package version 1.0.5. https://cran.r-project.org/package=BGLR

de los Campos, G., J. M. Hickey, R. Pong-Wong, H. D. Daetwyler, and M. P. Calus, 2012 Whole genome regression and prediction methods Applied to plant and animal breeding. Genetics 112.

Desta, Z. A., and R. Ortiz, 2014 Genomic selection: genome-wide prediction in plant improvement. Trends Plant Sci. 19: 592–601.

Dufresne, F., M. Stift, R. Vergilino, R. and B. K. Mable, 2014 Recent progress and challenges in population genetics of polyploid organisms: an overview of current state of theart molecular and statistical tools. Mol. Ecol. 23: 40–69.

Endelman, J. B., C. A. S. Carley, P. C. Bethke, J. J. Coombs, M. E. Clough et al., 2018 Genetic variance partitioning and genome-wide prediction with allele dosage information in autotetraploid potato. Genetics, 300685.

Ferrão, L. F. V., J. Benevenuto, I. de Bem Oliveira, C. Cellon, J. Olmstead et al., 2018 Insights into the genetic basis of blueberry fruit-related traits using diploid and polyploid models in a GWAS context. Front. Ecol. Evol. 6: 107.

Garcia, A. A., M. Mollinari, T. G. Marconi, O. Serang, R. R. Silva et al., 2013 SNP genotyping allows an in-depth characterisation of the genome of sugarcane and other complex autopolyploids. Sci. Rep. 3: 3399.

Garrison, E., and G. Marth, 2012 Haplotype-Based Variant Detection From Short-Read Sequencing. arXiv preprint arXiv:1207.3907.

Gorjanc, G., P. Bijma, and J. M. Hickey, 2015 Reliability of pedigree-based and genomic evaluations in selected populations. Genet. Sel. Evol. 47: 65.

Gouy, M., Y. Rousselle, D. Bastianelli, P. Lecomte, L. Bonnal et al., 2013 Experimental assessment of the accuracy of genomic selection in sugarcane. Theor. Appl. Genet. 126: 2575–2586.

Grandke, F., P. Singh, H. C. Heuven, J. R. De Haan, and D. Metzler, 2016 Advantages of continuous genotype values over genotype classes for GWAS in higher polyploids: a comparative study in hexaploid chrysanthemum. BMC Genomics 17: 672.

Guo, M., D. Davis, and J. A. Birchler, 1996 Dosage effects on gene expression in a maize ploidy series. Genetics 142: 1349–1355.

Guo, X., F. Cericola, D. Fè, M. G. Pedersen, I. Lenk, C. S. Jensen et al., 2018 Genomic Prediction in Tetraploid Ryegrass Using Allele Frequencies Based on Genotyping by Sequencing. Front. Plant. Sci. 9.

Gupta, V., A. D. Estrada, I. Blakley, R. Reid, K. Patel, M. D. Meyer et al., 2015 RNA-Seq analysis and annotation of a draft blueberry genome assembly identifies candidate genes involved in fruit ripening, biosynthesis of bioactive compounds, and stage-specific alternative splicing. Gigascience 4: 5.

Henderson, C. R., 1976 A simple method for computing the inverse of a numerator relationship matrix used in prediction of breeding values. Biometrics 32: 69–83.

Hieter, P., and T. Griffiths, 1999 Polyploidy--more is more or less. Science 285: 210–211.

Kerr, R. J., L. Li, B. Tier, G. W. Dutkowski, and T. A. McRae, 2012 Use of the numerator relationship matrix in genetic analysis of autopolyploid species. Theor. Appl. Genet. 124: 1271–1282.

Lee, W. P., M. P. Stromberg, A. Ward, C. Stewart, E. P. Garrison et al., 2014 MOSAIK: a hash-based algorithm for accurate next-generation sequencing short-read mapping. PloS one 9: e90581.

Lyrene, P. 2008 Breeding southern highbush blueberries. Plant Breed. Rev. 30: 353–414.

Lyrene, P., 2002 Development of highbush blueberry cultivars adapted to Florida. J. Am.Pomol. Soc. 56: 79.

Hayes, B. J., and M. E. Goddard, 2001 Prediction of total genetic value using genome-wide dense marker maps. Genetics, 157: 1819–1829.

Mollinari, M., and O. Serang, 2015 Quantitative SNP genotyping of polyploids with MassARRAY and other platforms, pp. 215–241.in Plant Genotyping edited by Humana Press, New York, NY.

Muir, W. M., 2007 Comparison of genomic and traditional BLUP estimated breeding value accuracy and selection response under alternative trait and genomic parameters. J. Anim. Breed. Genet. 124: 342–355.

Munoz, P. R., M. F. Resende, D. A. Huber, T. Quesada, M. D. Resende, D. B. Neal et al. 2014a. Genomic relationship matrix for correcting pedigree errors in breeding populations: impact on genetic parameters and genomic selection accuracy. Crop Sci. 54: 1115–1123.

Muñoz, P. R., M. F. Resende, S. A. Gezan, M. D. V. Resende, G. de los Campos et al., 2014b Unraveling additive from non-additive effects using genomic relationship matrices.Genetics 114.

Nyine, M., B. Uwimana, N. Blavet, E. Hřibová, H. Vanrespaille et al., 2018. Genomic prediction in a multiploid crop: genotype by environment interaction and allele dosage effects on predictive ability in banana. Plant Genome 11:170090.

Osborn, T. C., J. C. Pires, J. A. Birchler, D. L. Auger, Z. J. Chen et al., 2003 Understanding mechanisms of novel gene expression in polyploids. Trends Genet. 19: 141–147.

Pérez, P., and G. de Los Campos, 2014 Genome-wide regression ‖ prediction with the BGLR statistical package. Genetics 114.

R Core Team, 2018 R: A Language and Environment for Statistical Computing.

Resende, M. F. R., P. Munoz, J. J. Acosta, G. F. Peter, J. M. Davis et al., 2012 Accelerating the domestication of trees using genomic selection: accuracy of prediction models across ages and environments. New Phytol. 193: 617–624.

Resende, M. F., P. Muñoz, M. D. Resende, D. J. Garrick, L. R. Fernando et al., 2012 Accuracy of genomic selection methods in a standard dataset of loblolly pine (Pinus taeda L.). Genetics 111.

Resende, R. T., M. D. V. Resende, F. F. Silva, C. F. Azevedo, E. K. Takahashi et al., 2017 Assessing the expected response to genomic selection of individuals and families in Eucalyptus breeding with an additive-dominant model. Heredity 119: 245.

Schindelin, J., I. Arganda-Carreras, E. Frise, V. Kaynig, M. Longair et al., 2012 Fiji: an open-source platform for biological-image analysis. Nat. methods 9: 676.

Serang, O., M. Mollinari, and A. A. F. Garcia, 2012 Efficient exact maximum a posteriori computation for bayesian SNP genotyping in polyploids. PLoS One, 7: e30906.

Slater, A. T., N. O. Cogan, J. W. Forster, B. J. Hayes, and H. D. Daetwyler, 2016 Improving genetic gain with genomic selection in autotetraploid potato. Plant Genome, 9: 3.

Slater, A. T., G. M. Wilson, N. O. Cogan, J. W. Forster, and B. J. Hayes, 2014 Improving the analysis of low heritability complex traits for enhanced genetic gain in potato. Theor. Appl. Genet. 127: 809–820.

Soltis, D. E., and P. S. Soltis, 1999 Polyploidy: recurrent formation and genome evolution. Trends Ecol. Evol. 14: 348–352.

Soltis, D. E., P. S. Soltis, D. W. Schemske, J. F. Hancock, J. N. Thompson et al., 2007 Autopolyploidy in angiosperms: have we grossly underestimated the number of species?. Taxon, 56: 13–30.

Soltis, D. E., C. J. Visger, D. B. Marchant, and P. S. Soltis, 2016 Polyploidy: pitfalls and paths to a paradigm. Am. J. Bot. 103: 1146–1166.

Spoelhof, J. P., P. S. Soltis, and D. E. Soltis, 2017 Pure polyploidy: closing the gaps in autopolyploid research. J. Syst. Evol. 55: 340–352.

Sverrisdóttir, E., S. Byrne, E. H. R. Sundmark, H. Ø. Johnsen, H. G. Kirk et al., 2017 Genomic prediction of starch content and chipping quality in tetraploid potato using genotyping-by-sequencing. Theor. Appl. Genet. 130: 2091–2108.

Uitdewilligen, J. G., A. M. A. Wolters, B. Bjorn, T. J. Borm, R. G. Visser et al., 2013 A next-generation sequencing method for genotyping-by-sequencing of highly heterozygous autotetraploid potato. PloS one 8: e62355.

VanRaden, P. M., 2008 Efficient methods to compute genomic predictions. J. Dairy Sci. 91: 4414–4423.

Zhang, Z., X. Ding, J. Liu, D. J. de Koning, and Q. Zhang, 2011 Genomic selection for QTL-MAS data using a trait-specific relationship matrix. BMC Proc. 5: S15.

